# Bacterial Wastewater-Based Epidemiology Using Surface-Enhanced Raman Spectroscopy and Machine Learning

**DOI:** 10.1101/2024.07.22.604506

**Authors:** Liam K. Herndon, Yirui Zhang, Fareeha Safir, Babatunde Ogunlade, Halleh B. Balch, Alexandria B. Boehm, Jennifer A. Dionne

## Abstract

Wastewater-based epidemiology (WBE) is a powerful tool for monitoring community disease occurrence, but current methods for bacterial detection suffer from limited scalability, the need for *a priori* knowledge of the target organism, and the high degree of genetic similarity between different strains of the same species. Here, we show that surface-enhanced Raman spectroscopy (SERS) can be a scalable, label-free method for detection of bacteria in wastewater. We preferentially enhance Raman signal from bacteria in wastewater using positively-charged plasmonic gold nanorods (AuNRs) that electrostatically bind to the bacterial surface. Transmission cryoelectron microscopy (cryoEM) confirms that AuNRs bind selectively to bacteria in this wastewater matrix. We spike the bacterial species *Staphylococcus epidermidis, Staphylococcus aureus, Serratia marcescens*, and *Escerichia coli* and AuNRs into filter-sterilized wastewater, varying the AuNR concentration to achieve maximum signal across all pathogens. We then collect 540 spectra from each species, and train a machine learning (ML) model to identify bacterial species in wastewater. For bacterial concentrations of 10^9^ cells/mL, we achieve an accuracy exceeding 85%. We also demonstrate that this system is effective at environmentally-realistic bacterial concentrations, with a limit of bacterial detection of 10^4^ cells/mL. These results are a key first step toward a label-free, high-throughput platform for bacterial WBE.

---

Wastewater based epidemiology (WBE) is an important tool for surveilling infectious disease occurrence in a community.^[1]^ WBE has previously been used to monitor the spread of typhoid and polio;^[2, 3]^ within the last four years, its use has expanded globally for tracking COVID-19, MPOX, and other important diseases including measles, HIV, influenza, and RSV.^[4–9]^ Successful use cases of WBE mostly focus on viral diseases. Expanded use of WBE to monitor bacterial infections would greatly benefit public health. An estimated 7.7 million people died of bacterial infections in 2019, accounting for 13.6% of all deaths that year.^[10]^ Bacterial infections are currently the second leading cause of death worldwide, and this mortality rate is expected to increase with the rise of antimicrobial resistance.^[11]^ Recently, WBE has been extended for tracking selected bacterial (*Salmonella*) and fungal (*Candida auris*) infections,^[12, 13]^ as well as to track the emergence and spread of antimicrobial-resistant (AMR) bacteria within communities.^[14–16]^ The ability to broadly and accurately monitor bacterial pathogens at a population level and to address outbreaks before they rise to the level of pandemics could be critical to public health.

State-of-the-art WBE methods use nucleic-acid detection methods to quantify pathogen-specific nucleic-acids. The most commonly-used approach is digital polymerase chain reaction (PCR) owing to its high sensitivity and specificity, reduced potential for inhibition of the reactions, and high potential for multiplexing assays in the same reaction.^[17]^ However, PCR requires the identification of a unique, pathogen-specific nucleic-acid sequence for assay design. This requirement is challenging to scale to the many hundreds to thousands of possible bacterial species and strains and their associated antibiotic susceptibility. Further, virulence of a pathogen can be determined by phenotypic signatures (such as protein expression levels and post-translational modifications) which are not accessible from genomic or transcriptomic analysis alone.^[18]^ Finally, *a priori* knowledge of pathogen genomic sequences also means that new, emerging antibiotic resistant strains could be challenging to identify with PCR.

Vibrational spectroscopy, including Raman and infrared (IR) spectroscopy, offers a label-free route towards sensitive, specific, and scalable identification of broad ranges of pathogenic bacteria in wastewater. Because these techniques directly measure molecular vibrations, they do not require targeted probes or *a priori* knowledge of the target species.^[19]^ Raman spectroscopy, which uses a monochromatic light source to measure vibrational energy levels based on the redshift of inelastically-scattered photons, is particularly suitable for wastewater analysis. Raman spectroscopy experiences less absorption in aqueous samples when compared to IR spectroscopy. Raman spectroscopy is also generally more field-deployable and less expensive, owing to its visible excitation and detection wavelengths.^[20, 21]^ Different species and strains of bacteria have been shown to possess unique Raman “fingerprints”, allowing Raman identification of clinically-relevant information at the species and subspecies level.^[22]^ Based on our and others’ libraries of pathogen Raman spectra in water, sputum, blood, and other clinical samples, we propose that Raman can be used for bacterial WBE by measuring light scattered off wastewater in a liquid well (Fig. 1a).^[22–26]^

**Figure 1.**
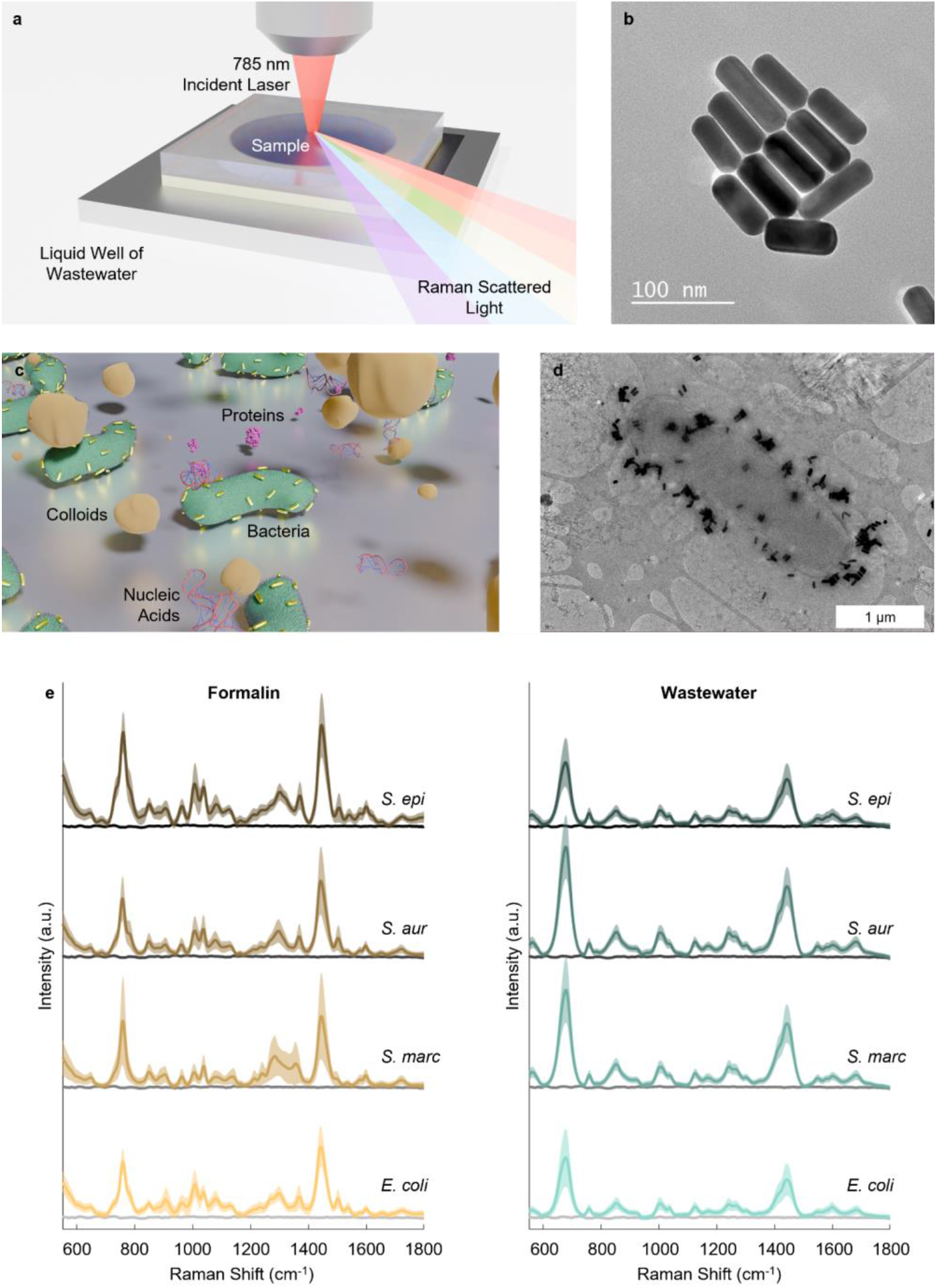
(a) Raman spectra are collected from liquid samples in microfluidic wells. (b) Electron micrograph of AuNRs used for SERS. (c) In the liquid wells, AuNRs selectively enhance scattered signals from bacteria over a complex mixture of background components, such as colloids, proteins, and nucleic acids. (d) Cryoelectron micrograph of electrostatic binding of AuNRs to *E. coli* surface. (e) SERS spectra of four model species collected in formalin and wastewater. Grey lines indicate negative control without AuNRs. SERS spectra are averaged over a minimum of 216 samples and unenhanced spectra are averaged over a minimum of 36 samples. Shaded regions indicate 1 standard deviation error

Machine learning (ML) techniques can be used to interpret bacterial Raman spectra. Several previous studies have used ML models including random forest classifiers and neural networks to identify bacterial species from Raman spectra, achieving high accuracies (currently up to >99%) across many dozens of pathogens.^[22, 25, 27, 28]^ ML has been used to identify clinically relevant properties of bacteria at the subspecies level, including antibiotic susceptibility.^[22, 24, 29, 30]^ Additionally, ML techniques have been used to identify spectral features that account for differences between species, providing insight into the biochemical pathways and processes underlying Raman classification.^[25]^

Combined with ML, surface-enhanced Raman spectroscopy (SERS) with plasmonic gold nanorods (AuNRs) has significant potential to identify pathogens in complex samples. In this method, the AuNRs focus the incident field onto the analytes, with reported enhancement factors ranging from 10^3^-10^9^.^[31–33]^ This technique enables rapid, sensitive characterization of analytes through Raman spectroscopy.^[34–39]^ For example, in wastewater, Liu et al. developed a SERS system to measure estrogen concentrations, wherein Raman signal from a secondary reporter increased in wastewaters with higher estrogen concentrations.^[40]^ To achieve *reporter-free* detection of bacteria in wastewater, we utilize the electrostatic interaction between bacteria (which are generally negatively-charged) and AuNRs (which can be synthesized with positively-charged ligands such as cetrimonium bromide (CTAB)) (Fig. 1b-c).

We have previously shown in deionized (DI) water and in blood that electrostatic interactions will allow AuNRs to bind selectively to the cell surface of almost any bacteria (including gram positive and gram negative species), to enhance bacterial Raman over the background medium’s Raman signature.^[23, 25]^ We hypothesize that even in a sample as complex as wastewater, which contains a mixture of a variety of biomolecules and waste products, AuNRs can enhance bacterial Raman signal over the background, enabling label-free identification of bacteria in wastewater (Fig. 1d).

Here, we establish the feasibility of WBE using SERS and ML. As a proof of concept, we consider four bacterial species, *Staphylococcus epidermidis* (*S. epidermidis*), *Staphylococcus aureus* (*S. aureus*), *Serratia marcescens* (*S. marcescens*), and *Escherichia coli* (*E. coli*) spiked into filter-sterilized wastewater with CTAB-functionalized AuNRs, and demonstrate SERS from this spiked wastewater with an acquisition time of only 10 seconds. Additionally, we use transmission cryoelectron microscopy (cryoEM) to confirm AuNR binding to the cell surface through strong electrostatic interactions. We investigate the relationship between AuNR concentration and bacterial Raman signal in wastewater, and demonstrate that the AuNRs have a primarily enhancing, rather than quenching effect among the tested concentrations. Additionally, we show that bacterial SERS spectra collected in wastewater can be used to accurately predict bacterial species: we train a convolutional neural network (CNN) to classify them with >85% accuracy. Finally, we demonstrate that this method can detect the presence of bacteria in wastewater at environmentally-relevant concentrations as low as 10^4^ cells/mL.

## Results and Discussion

### SERS Enhancement of Bacterial Signal in Wastewater

AuNRs were synthesized using a seed growth method using CTAB.^[41]^ The cylindricity of CTAB micelles gives the nanoparticles a rod shape, and the trimethylammonium group on the ligand gives the AuNRs a positive surface charge, which we have previously shown to enable electrostatic interactions with negative charges on the bacterial surface.^[23]^

AuNRs were designed to have a longitudinal surface plasmon resonance (LSPR) at approximately 700 nm and dimensions of approximately 75 nm by 30 nm, to optimize the balance between enhancement and quenching for SERS with a 785 nm laser (Fig. 1b).^[23, 42, 43]^ All AuNRs used for these experiments had resonances of 700-720 nm, lengths of 73.5-79.4 nm, and widths of 28.1-29.1 nm. Moreover, the two AuNRs batches used for SERS experiments had resonances of 700 nm, lengths of 75.3 nm and 73.5 nm, and widths of 28.1 nm and 29.1 nm (Fig. S1).

We used clinically-relevant pathogens *S. epidermidis, S. aureus, S. marcescens*, and *E. coli* as model organisms to determine the effects of wastewater on AuNR enhancement of bacterial Raman spectra. *S. marcescens, S. aureus*, and *S. epidermidis* have respective surface charge densities 4, 29, and 102 times greater than *E. coli* and have stronger electrostatic binding to AuNRs in deionized water.^[23, 44]^

Here we tested bacterial SERS in a wastewater matrix, which was prepared from the liquid portion of wastewater from a local wastewater treatment plant. We filter-sterilized the wastewater through a 0.22 μm pore-size filter to remove endogenous bacteria, in addition to other cells and particles, while preserving the colloids, dissolved ions, biomolecules, and various dissolved molecules in wastewater. Samples were then formalin treated for biosafety. For each experiment, this wastewater matrix was spiked with one of the model species and mixed 1:1 with AuNRs for a final concentration of 150 μg/mL AuNRs and 10^9^ cells/mL, which is approximately equal to the total bacterial concentration in untreated wastewater.^[45, 46]^ As a control, samples were also prepared in formalin-treated DI water.

Samples were transferred to a liquid well for Raman analysis (Fig. 1a). Raman spectra were collected from each species of bacteria with and without AuNRs in formalin and filter-sterilized wastewater, acquired over 10 seconds with a 785 nm laser at a power of 11 mW and a laser spot size of 2 μm. A minimum of 216 spectra per species werecollected with AuNRs (and 36 spectra were collected without AuNRs). In samples without AuNRs, no bacterial Raman peaks were observed (Fig. 1e), consistent with our previous liquid Raman results with similarly low exposure times and power densities.^[23]^ In samples with cells and AuNRs in formalin, strong Raman peaks are visible (Fig. 1e).These peaks overlap strongly with dozens of wavenumbers known to be associated with biomolecules on the bacterial surface. These features include particularly strong peaks at 760 cm^-1^, which is associated with adenine; 851 cm^-1^, which is associated with thymine; 1,040 cm^-1^, which is associated with aryl; 1,125 cm^-1^, which is associated with phosphate; and 1,599 cm^-1^, which is associated with carboxyl.^[25]^ All of these major peaks are conserved across the four species, which is expected: differences between the Raman spectra of different bacterial species tend to be subtle and difficult to discern by eye.^[22]^

In samples with bacteria and AuNRs in a filtered wastewater matrix, strong peaks are still present. There are, however, some differences between these spectra and those collected in formalin. The most obvious difference is a peak at 670 cm^-1^ that is only present in bacterial spectra collected in wastewater. This band is characteristic of the C-S stretching vibration of the thiol group in cysteine, which is known to covalently bond to noble metal nanoparticles. We therefore hypothesize that this peak is the result of fecal proteins bonding to the AuNRs through their cysteine residues. Despite this new thiol peak, spectra collected in wastewater are otherwise qualitatively similar to those collected in formalin. All major peaks are conserved, including the biological peaks at 760 cm^-1^, 851 cm^-1^, 1,040 cm^-1^, 1,125 cm^-1^, and 1,599 cm^-1^ that were previously highlighted.

Despite these qualitative similarities, the spectra collected in filtered wastewater are not identical to those collected in formalin; the intensities of individual peaks vary between the two matrices. Some, but not all, of these peaks decrease in intensity in wastewater compared to formalin. The greatest decrease was observed for the 760 cm^-1^ peak, which decreases by 76.0-88.4% depending on species. The 1,040 cm^-1^ peak also decreases across all species by 51.0-77.8%. The 851 cm^-1^, 1,125 cm^-1^, and 1,599 cm^-1^ peaks, however, do not display a statistically-significant change in intensity in formalin compared to wastewater, indicating that wastewater does not strongly affect the intensities of these peaks (Fig. S2).Thus, the complex environment of wastewater does not inherently decrease SERS enhancement of bacterial surface molecules. While it has a weakening effect with some compounds, it does not affect the enhancement of others. Moreover, the peaks that do decrease in intensity are still clearly observable. Overall, SERS is achievable from bacteria in a wastewater matrix.

### Characterization of Nanorod Binding to Selected Bacterial Species

CryoEM was used to further validate that bacterial Raman signal in wastewater was the result of AuNR binding to cells in this matrix. Samples were drop-cast onto glow-discharged lacey carbon grids, plunge frozen, then imaged using a transmission cryoEM. In samples in formalin, strong AuNR binding to bacteria was observed. Across all four model species, multiple dozens of AuNRs are bound to the surface of the average cell (Figs 2a,c,e,g). In formalin-treated wastewater, cryoEM images still show AuNR binding to all four species (Figs. 2b,d,f,h). For *S. epidermidis* and *E. coli*, the wastewater matrix does not decrease binding at all: the typical cell in wastewater has at least as many AuNRs bound to it as the typical cell in formalin (Figs. 2a,b,g,h, S3). For *S. aureus* and *S. marcescens*, a decrease in AuNR binding was observed in wastewater compared to formalin, but it does not entirely eliminate binding. In the case of *S. aureus*, tens of AuNRs are still bound to the surface of the typical cell (Figs. 2c,d, S3). And while *S. marcescens* has the greatest decrease in binding in wastewater relative to formalin, the typical cell still has multiple AuNRs bound to it (Figs. 2e,f, S3).

**Figure 2.**
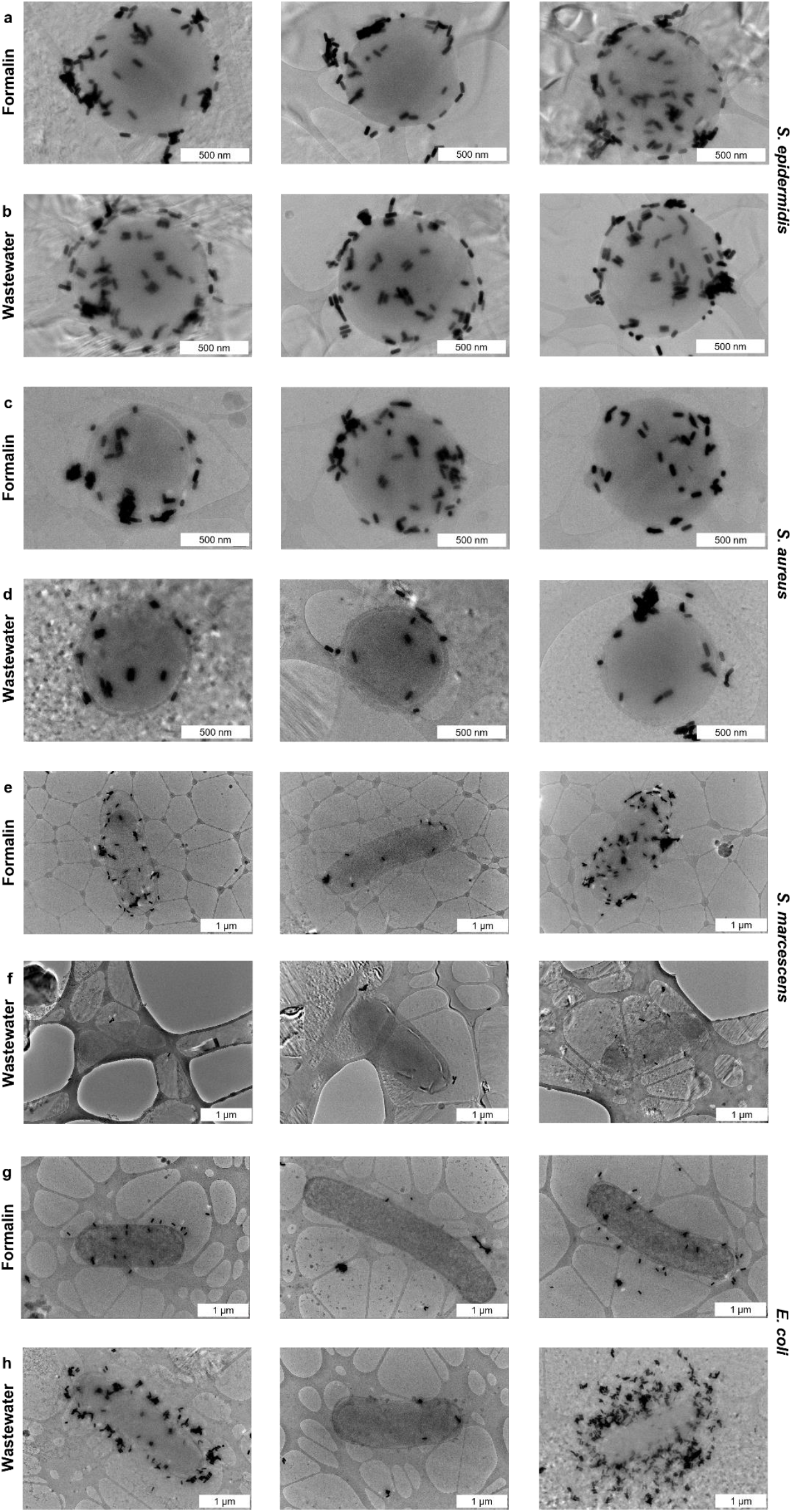
Cryoelectron micrographs of AuNRs bound to (a) *S. epidermidis* in formalin, (b) *S. epidermidis* in wastewater, (c) *S. aureus* in formalin, (d) *S. aureus* in wastewater, (e) *S. marcescens* in formalin, (f) *S. marcescens* in wastewater, (g) *E. coli* in formalin, and (h) *E. coli* in wastewater.

*S. epidermidis* has greater surface charge density than *S. aureus*, which, in turn, has greater surface charge density than *S. marcescens*. Thus, for these three species, we observed a relationship in which lower charge density leads to a greater decrease in AuNR binding in a wastewater matrix. This result is likely because weaker electrostatic interactions can be more easily disrupted by changes in the ionic strength of their environment. Interestingly, *E. coli* has the lowest surface charge density of the four species, but does not display any decrease in AuNR binding in wastewater compared to formalin. We therefore hypothesize that wastewater is a more favorable environment for AuNR binding in cases where the electrostatic interaction is very weak.

Thus, wastewater can decrease AuNR binding to the cell surface, but AuNR binding is never eliminated. Bacterial SERS has previously been achieved from samples with very few AuNRs directly bound to the cell surface, indicating that even low amounts of AuNR binding can produce SERS.^[23]^ It can therefore be concluded that AuNR binding to all four species in wastewater generates the enhanced Raman signal we observe from bacteria in wastewater.

### Assessment of Plasmonic Nanoparticle Concentration: Enhancement vs. Quenching

Due to the inherently absorbing nature of plasmonic nanoparticles, excess substrate can be detrimental to SERS, quenching the incident field more strongly than it enhances it. Therefore, a threshold nanoparticle concentration can exist, above which signal decreases.^[43]^ To ensure that the 150 μg/mL AuNR concentration used in this system is below this threshold, bacterial samples of each selected species were prepared in filtered wastewater with AuNRs at concentrations of 10 μg/mL, 50 μg/mL, and 100 μg/mL, in addition to the previous spectra collected at 150 μg/mL. A minimum of 144 Raman spectra were collected from each species at each concentration, and the mean intensities of the 760 cm^-1^, 851 cm^-1^, 1,040 cm^-1^, 1,125 cm^-1^, and 1,599 cm^-1^ peaks (which respectively correspond to adenine, thymine, aryl, phosphate, and carboxyl) for each species were compared (Fig. 3).

**Figure 3.**
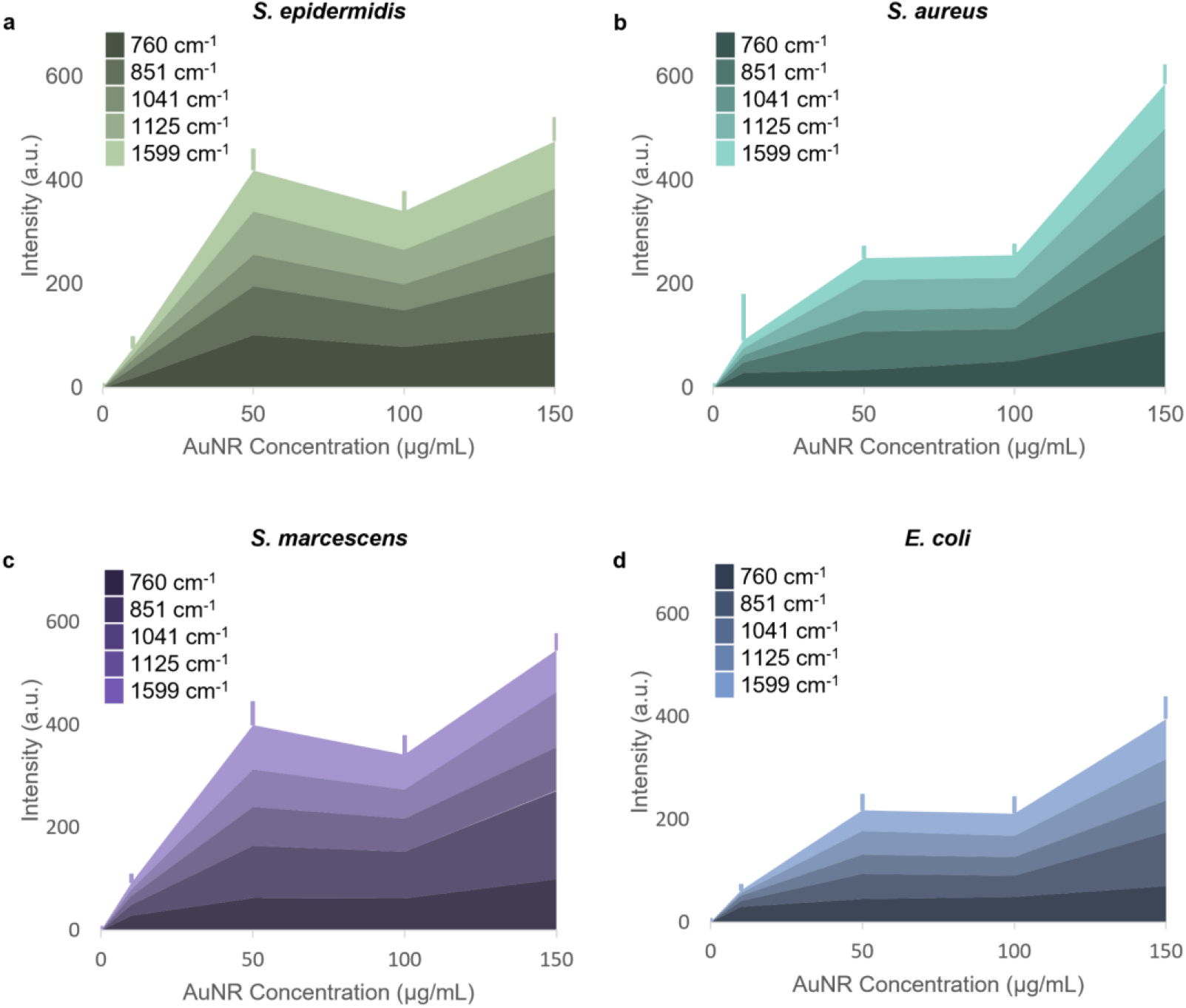
Stacked area chart depicting the intensities of selected bacterial peaks in wastewater depending on AuNR concentration. Each colored band indicates the intensity of one peak, and the total height of the chart indicates the sum of all five peak intensities. Error bars indicate standard deviation of 1599 cm^-1^ peak, which is representative of standard deviation for all peaks.

Across all species, the greatest signal was generally observed at 150 μg/mL. We note that there is a local maximum at AuNR concentrations of 50 μg/mL, compared to concentrations of 100 μg/mL. In these intermediate concentration regimes, it is possible that quenching outcompetes enhancement. Alternatively, it could be that nanorod electrostatic interactions are more stable with certain relative concentrations of bacteria and AuNRs. It is also possible that AuNRs are more likely to aggregate in certain concentration regimes, which can decrease enhancement. Overall, we determine that a concentration of 150 μg/mL AuNRs enhances Raman signal more than it quenches it, making it suitable for bacterial SERS in wastewater systems.

### Machine Learning Classification of Bacteria

As numerous biological molecules present Raman peaks in the fingerprint region, we applied ML to classify the Raman spectra and assess the utility of SERS for bacterial identification in wastewater. 540 spectra from each bacterial species were collected in wastewater with 150 μg/mL AuNRs and analyzed using a one-dimensional CNN consisting of a convolutional layer followed by seven fully-connected residual layers. The CNN can account for peak shape and has previously successfully classified bacterial Raman spectra.^[22]^

This model was trained on the bacterial Raman spectra collected in wastewater from 725 cm^-1^ to 1800 cm^-1^. This range was selected because it excludes the non-bacterial 670 cm^-1^ thiol peak from wastewater, which could lead to overfitting. The model was validated using k-folds cross-validation. In this process, the spectra from each species were split 4:1 into a training set and a test set. The model was then trained on the training set, and validated using the test set. This process was repeated using 500 different train-test splits. This model identified *S. epidermidis* with 86.4% accuracy, *S. aureus* with 77.0% accuracy, *S. marcescens* with 86.5% accuracy, and *E. coli* with 91.0% accuracy (Fig. 4a). As CNNs are typically trained using, at minimum, several thousand objects per class in the training set, which is far greater than the 540 here, we expect that this accuracy will increase in future research with larger datasets. Additionally, overall accuracy is decreased by an *S. aureus* classification accuracy that is markedly lower than that for the other organisms. A majority of this *S. aureus* misclassification is the result of the model misidentifying this organism as *S. marcescens*. Thus, much of this model’s inaccuracy stems from misclassification between *S. aureus* and *S. marcescens*, which we hypothesize can be overcome by generating a larger dataset of spectra from those two specific species.

**Figure 4.**
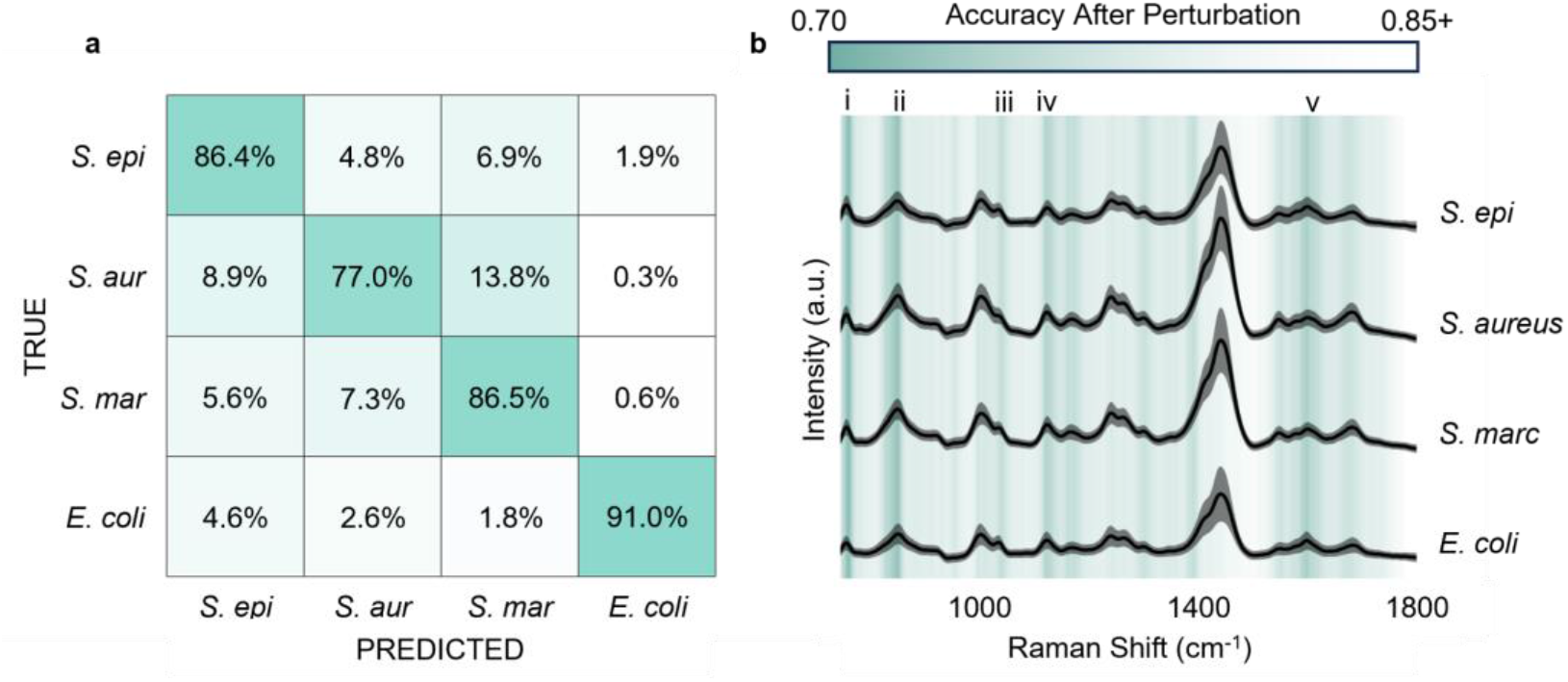
(a) Confusion matrix showing >85% accuracy of bacterial classification using a CNN trained on bacterial SERS spectra collected in wastewater. (b) Graph of accuracy of CNN when the test set is perturbed at various wavenumbers. Sharp decreases can be seen with perturbation at bands associated with (i) adenine, (ii) thymine, (iii) aryl, (iv) phosphate, and (v) carboxyl.

To ensure that this accurate ML classification was based on biologically relevant Raman peaks, relevant wavenumbers were identified using an iterative perturbation method. For each train-test split, all spectra in the test set would be perturbed by adding an artificial peak at a specific wavenumber. The effect on classification accuracy of perturbation at this wavenumber would be recorded. This process was iteratively repeated for all wavenumbers in the spectra. Perturbation of a wavenumber relevant to classification can be expected to cause a strong decrease in accuracy, so this method can identify these relevant wavenumbers.^[25]^ We observed that wavenumbers relevant to classification strongly overlapped with known bacterial Raman peaks. For example, perturbation at 760 cm^-1^ (which corresponds to adenine) decreases accuracy by 11.8%, perturbation at 851 cm^-1^ (which corresponds to thymine) decreases accuracy by 11.2%, perturbation at 1,040 cm^-1^ (which corresponds to aryl) decrease accuracy by 6.4%, perturbation at 1,125 cm^-1^ (which corresponds to phosphate) decreases accuracy by 8.0%, and perturbation at 1,599 cm^-1^ (which corresponds to carboxyl) decreases accuracy by 9.2% (Fig. 4b). Thus, the >85% classification accuracy is based on differences between the Raman spectra of different bacterial species in wastewater stemming from the different chemical compositions of these species.

### Limit of Bacterial Detection

The concentrations of clinically relevant species in raw wastewater span many orders of magnitude. While the total concentration of bacteria in wastewater has been reported on the order of 10^9^ cells/mL, the concentrations of individual species tend to be far lower.^[45]^ *E. coli*, for example, has reported concentrations ranging from the order of 10^2^-10^5^ cells/mL.^[47]^ To identify individual bacterial species, it is therefore necessary to be able to detect bacteria at concentrations several orders of magnitude below 10^9^ cells/mL. To assess the limit of detection (LoD) for bacterial SERS in wastewater, samples were prepared in a wastewater matrix with 150 μg/mL AuNRs and bacteria at concentrations of 0 cells/mL, 10^4^ cells/mL, 10^5^ cells/mL, 10^6^ cells/mL, 10^7^ cells/mL, 10^8^ cells/mL, and 10^9^ cells/mL. At each concentration, 756 Raman spectra were collected, including a minimum of 108 spectra from each species (Fig. 5a). At these lower concentrations, biological peaks are still present in the spectra. This result is likely the result of the wastewater matrix containing biomolecules similar to those on the bacterial cell surface. However, the intensities of many of these peaks decrease at low bacterial concentration, indicating a strong contribution to these peaks from the bacterial surface (Fig. S4).

**Figure 5.**
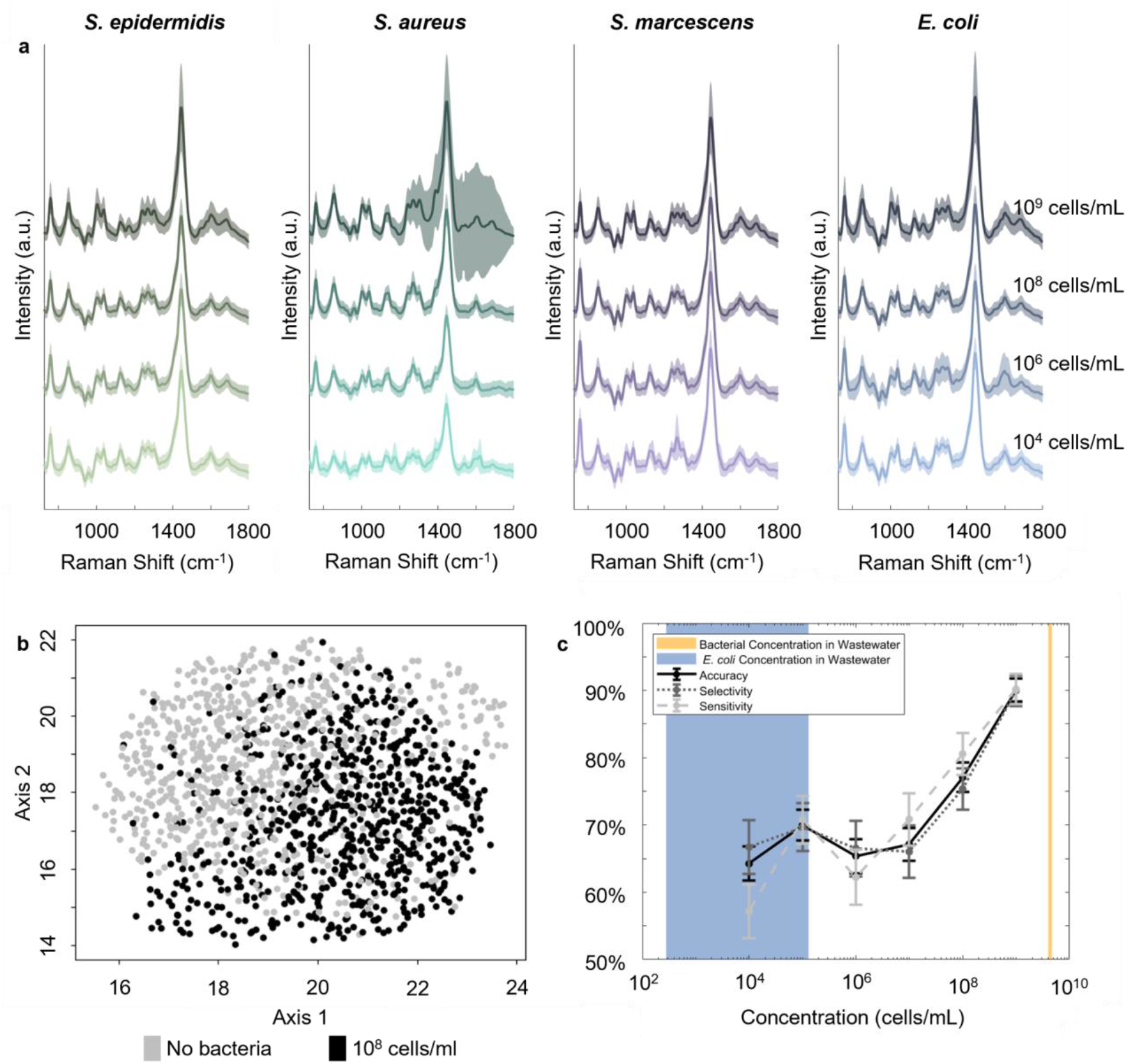
(a) SERS spectra of four model species collected in wastewater at varying bacterial concentrations. (b) UMAP showing clustering of wells without bacteria vs wells with 10^8^ cells/mL. (c) Sensitivity, selectivity, and accuracy of KNN classifiers trained to detect presence of bacteria at varying concentrations. Shaded regions indicate concentrations reported in literature for total bacteria in wastewater and *E. coli* in wastewater.

To test the limit of detection, ML techniques were used to classify between spectra from samples containing bacteria at each concentration, and samples without bacteria. For example, Fig. 5b shows a uniform manifold approximation and projection (UMAP) plotting a two-dimensional projection of a set of spectra with bacteria at 10^8^ cells/mL, and a set of spectra containing no bacteria. While this projection does not show wholly separate clusters, spectra with and without bacteria are generally localized to different regions of the two-dimensional vector space, indicating noticeable differences between spectra from wells with and without bacteria at this concentration.

For a more robust comparison of spectra from samples with bacteria at each concentration against spectra with no bacteria, a k-nearest neighbors (KNN) classifier was used. In this process, principal component analysis (PCA) was performed on a dataset containing all the spectra with 10^9^ cells/mL and all the spectra with no bacteria. The top 3 eigenvectors from this PCA were then used to project all of the spectra at all bacterial concentrations into 3 dimensions while preserving important information. A KNN classifier was then trained on the projected spectra at each concentration to classify between spectra from wells containing that concentration of bacteria, and wells containing no bacteria. The accuracy of each model was tested across 500 train-test splits using the same methods used to validate the CNN.

At concentrations of 10^4^-10^7^ cells/mL, sensitivity, selectivity, and accuracy are approximately constant, all falling in the 55-70% range. At 10^8^ cells/mL, sensitivity, selectivity, and accuracy respectively increase to 80.6±3.1%, 75.4±3.4%, and 77.1±2.2%. At 10^9^ cells/mL, sensitivity, selectivity, and accuracy respectively reach 90.2±2.3%, 90.0±2.4%, and 90.1±1.7%. (Fig. 5c). The threshold concentration is around 10^7^ cells/mL, above which accuracy increases significantly (Fig. S5). This threshold matches the concentration above which a dense layer of cells forms on the bottom of the well Fig. S6). Above the threshold concentration, we hypothesize that cells are present in the laser spot for most spectral acquisitions, so the spectra contain information on the cell surface itself; below the threshold, molecules are exchanged between the cells and the surrounding medium to give rise to Raman peaks.

It is notable that the presence of bacteria could be detected with >50% accuracy, even at low concentrations at which bacteria were not in the laser spot. This result indicates that bacteria release enough molecules into the wastewater matrix to be detected by Raman. There is therefore potential for future Raman-based bacterial detection systems that do not require direct irradiation of the cells themselves. Together, these results suggest that SERS is capable of detecting bacteria at environmentally-relevant concentrations. We note, however, that for this system to be implemented in a real wastewater system, we will need methods to identify signals from individual species in raw wastewater rather than filter-sterilized wastewater; the former will contain broad classes of bacteria, as well as protozoa and other organic and inorganic particles.

## Conclusions

We have established that bacterial SERS can be achieved in a filter-sterilized wastewater matrix, en-route to bacterial WBE. We demonstrate that bacterial SERS in wastewater generates Raman peaks comparable to those observed in formalin. Additionally, cryoEM images show that AuNRs bind to the bacterial surface in this complex matrix, indicating that the various dissolved substances in wastewater do not adversely affect this interaction, and indicating that SERS enhancements in wastewater are the result of a bacteria-AuNR interaction. We demonstrate that a high AuNR concentration of 150 μg/mL is suitable for SERS, enhancing signal rather than quenching it. We also demonstrate that bacterial SERS spectra collected in filtered wastewater are sufficiently robust for species identification: an ML model trained on spectra from four clinically relevant bacterial species in wastewater could distinguish between these species with >85% accuracy. Combined with ML, this platform can detect bacteria in wastewater at concentrations as low as 10^4^ cells/mL.

Our proof-of-concept work marks a critical first step toward scalable wastewater monitoring of bacterial pathogens. Future work remains to bring this technology to practice. First, it would be crucial to expand the CNN training dataset to include a larger number of species and tens of thousands of spectra per species. This expanded dataset would enable SERS monitoring of a fuller breadth of the bacterial pathogens present in wastewater and have sufficient data per class for optimal CNN performance.

Additionally, work is needed to diversify the wastewater background training data (eg, from a variety of geographic locations), as well as the bacterial Raman cataloging. Future datasets would ideally contain various strains of each species cultured under a range of conditions and spiked into wastewater from various sources. There is also potential for synthetic data augmentation techniques to diversify data. Such diversity would prevent potential overfitting to a specific set of conditions.

Improvements in identification accuracy could be further improved with a platform to collect spectra from few-to-single cells in wastewater, allowing characterization of wastewater on a cell-by-cell basis. In recent years, systems have emerged integrating bioprinting and microfluidics with Raman spectroscopy, which have potential for single cell isolation.^[25, 48]^ Additionally, there is promise for enrichment techniques, such as dielectrophoresis, for concentrating bacteria near electrodes, to maximize their Raman signal, even at low bacterial concentrations. We anticipate that a Raman system that combines bioprinting or microfluidics with enrichments could be powerful as a means of sensitively generating single-cell spectra.

In summary, SERS has significant potential as a tool for wastewater monitoring, and these results lay the groundwork for the implementation of this epidemiological tool. In the future, we hope that this rapid, label-free technique will allow bacterial outbreaks to be detected and addressed before they rise to the level of crises.

## Methods

### Wastewater preparation

Raw, untreated wastewater influent was collected on 19 April 2022 as an eight-hour composite sample using a composite automated sampler from a large wastewater treatment plant located in the Bay Area of California, USA. This wastewater sample is likely representative of wastewater from large urban areas in the USA. Wastewater was stored for a period of several weeks at 4°C, during which solid components separated from liquid components through gravitational setting. The liquid portion of this wastewater was recovered using a sterile pipetter and subsequently filtered through a 0.22 μm pore size syringe filter (09-720-3, Fisher) to remove particles including bacteria and protozoa. Filter-sterilized wastewater was then mixed 1:1 by volume with 10% formalin for biosafety. Filtered, formalin-treated wastewater samples were stored at 4°C for up to two years.

### Nanorod synthesis and Characterization

To achieve our target LSPR and AuNR width, AuNRs were synthesized using a modified version of the recipe described in subfigure 3a of the 2013 manuscript by Ye *et al*.^[41]^ The scale of this synthesis was decreased by a factor of 10 from the scale described in the manuscript, and hydrochloric acid and seed concentrations were sometimes modified to correct for effects of seasonal changes in humidity on AuNR growth. The nanorods were washed once, which is sufficient to prevent cytotoxicity from CTAB, while leaving sufficient CTAB on the AuNR surface to maintain a positive surface charge and prevent aggregation.^[23]^ AuNRs were washed by centrifuging the synthesis for 20 minutes at 5,400 g, removing the supernatant, resuspending in 50 mL DI water, centrifuging a second time with the same parameters, removing supernatant, and resuspending in 5 mL DI water. AuNR resonance was measured by collecting their extinction spectrum from 400-900 nm using a Cary 6000i UV-Vis-NIR spectrometer. AuNR concentration was determined by using a Thermo Scientific ICAP 6300 Duo View Optical Emission Spectrometer to measure the concentration of gold in a sample containing the washed AuNRs diluted 1:200 in 5% nitric acid. To analyze individual AuNRs, images of fifty AuNRs were recorded with an FEI Tecnai G2 F20 X-TWIN Transmission Electron Microscope. Each AuNR in these images was measured using the line tool in ImageJ.

Three batches of AuNRs were synthesized for the experiments discussed in this paper. The AuNRs for the experiments in Figs. 1, 3, and 4 had an LSPR at 700 nm, an average length of 75.3 nm with 9.6% standard deviation, and an average width of 28.1 nm with 8.9% standard deviation. The AuNRs for the experiments in Fig. 2 had an LSPR at 720 nm, an average length of 79.4 nm with 10.8% standard deviation, and an average width of 28.9 nm with 9.0% standard deviation. The AuNRs for the experiments in Fig. 5 had an LSPR at 700 nm, an average length of 73.5 nm with 21.5% standard deviation, and an average width of 29.1 nm with 24.1% standard deviation.

### Cell Culture

*S. epidermidis* (ATCC 12228), *S. aureus* (which was a clinical sample obtained from the Stanford Hospital), *S. marcescens* (ATCC 13880), and *E. coli* (ATCC 25922) were cultured on Trypticase Soy Agar 5% Sheep Blood plates (221239, BD) at 37°C for 16 hours. Colonies from these plates were then cultured in 12.5 mL Lysogeny broth culture medium (10-855-021, Fisher) in 50 mL bio-reaction tubes (229476, CellTreat) at 37°C with 120 rpm shaking for 16 hours. All cultures were incubated in a Thermo Scientific MaxQ 4450 incubator. Cultures were stored at 4°C, and experiments were not performed with solid cultures more than 14 days old or liquid cultures more than 5 days old.

### Preparation of Cell/AuNR Mixtures

Cell concentration was measured by diluting an aliquot of liquid culture 1:1,000 and counting on a Bright-Line Hemacytometer. Cell aliquots were washed 3 times by centrifuging for 4 minutes in a mySPINTM 6 Mini Centrifuge, removing supernatant and resuspending in DI water between centrifugations. After the washes, the cells were resuspended in either 5% formalin or filtered and formalin-treated wastewater at double their desired concentration. Additionally, aliquots of AuNRs were diluted in DI water to double the desired final concentration. A cell aliquot and an AuNR aliquot were then mixed in a 1:1 ratio by volume.

### Liquid Well Preparation

A 0.5 mm silicon chip and a 0.5 mm JGS2 grade, double-side polished fused silica chip (WA1001, MSE Supplies) were plasma cleaned for 5 minutes at 100 W with an 85 mTorr base pressure and a 2 SCCM oxygen flow rate in a March Instruments PX-250 Plasma Asher to decrease hydrophobicity and remove residual adhesive from previous experiments. 4 layers of double-sided scotch tape (6137H-2PC-MP, 3M) were stacked on each other for a final thickness of approximately 0.1 mm,^[32]^ and a hole was punched in the middle of this stack using a Bostitch Office EZ Squeeze One-Hole Punch. This tape was then placed on the plasma-cleaned silicon. A 10 μL sample was pipetted into the hole in the tape, and the top was sealed with the plasma-cleaned silica. After experiments, wells were disassembled by dissolving tape adhesive in 10% isopropyl alcohol (IPA). Silicon and silica substrates were then wiped down with IPA and reused in subsequent experiments.

### CryoEM

To prepare cryoEM samples, 300 mesh molybdenum lacey carbon grids (LC300-MO, Electron Microscopy Sciences) were glow discharged for 10 seconds in a PELCO easiGlow Glow Discharge Cleaning System at 15 mA and 0.40 mbar. 3 μL of cell/AuNR mixture was subsequently drop-cast onto each grid, which was then back-blotted for 3 seconds and frozen using a Vitrobot Mark IV automatic plunge freezer, with blot force set to 3. Cells were imaged using a Glacios 2 Cryo-Transmission Electron Microscope.

### Raman Spectroscopy

Raman spectra were collected using a Horiba Xplora+ Confocal Raman microscope with a 600 gr/mm grating, a 300 μm pinhole, and a 785 nm laser with 11 mW power output. Spectra were collected through an Olympus Plan N 10x objective (N1215800, Olympus), with a spot size of approximately 2 μm. Each spectrum was collected through two 5-second acquisitions. Spectra were collected from 550-1800 cm^-1^, with a multiple accumulation spike filter and an intensity correction system (ICS). Spectra were collected in batches of 36 by collecting a 6×6 map over a 200×200 μm area. Between 1 and 3 maps were collected from each well. For each species and concentration, Raman was performed on a minimum of two wells containing cells from separate liquid cultures, aside from negative controls without AuNRs and *S. aureus* samples in Figure 5, which were collected from a single well for each sample.

### Data Processing

Spectra were processed using Python. Each spectrum was truncated to only include wavenumbers from 550-1800 cm^-1^. To avoid overfitting to the 670 cm^-1^ thiol peak, whose intensity varied significantly between spectra, spectra being processed for ML were truncated to a range of 725-1800 cm^-1^. To remove the fluorescent background, a fifth-order polynomial fit was subtracted from each spectrum using peakutils.baseline. To remove background from silicon in the wells, the baseline-subtracted spectrum of a well containing DI water was subtracted from each baseline-subtracted spectrum. For normalization, each spectrum then had its mean value subtracted and then was divided by its standard deviation, to achieve a mean of 0 and a standard deviation of 1.

### Machine Learning

All ML was performed using Python. PCA was performed on processed and normalized spectra using sklearn.decomposition.PCA. UMAP was performed using umap.UMAP with min_dist=1 and n_neighbors=1500. The CNN was built using pytorch_lightning with the architecture and parameters we previously described in the 2019 manuscript by Ho *et al*.^[22]^ The KNN classifier used sklearn.neighbors. KNeighborsClassifier, with n_neighbors=40. Each classifier was validated using k-folds cross validation, with each class randomly divided into five folds using sklearn.model_selection.KFold. The model iterated through the five folds, using each one as the test set, while using the other four as the training set. This process was repeated with a new set of folds 100 times, for a total of 500 training and test sets.

For the CNN, iterative perturbation was performed for each of the 500 test sets. This process was repeated for all wavenumbers in the spectra. For each spectrum, a Lorentzian curve was generated using the equation: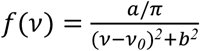, where *ν* is a wavenumber in cm^-1^, *ν*_*0*_ is the perturbed wavenumber in cm^-1^, *a* is a value randomly selected from an even distribution from 0-250 using random.random, and b=5. A Lorentzian centered at a specific wavenumber would be added to a spectrum to perturb it at that wavenumber. For each of the 500 train-test splits, each spectrum in the test set was perturbed at each wavenumber.

## Acknowledgements

This work was funded with grants from the Chan Zuckerberg Biohub, San Francisco; and Stanford Woods Institute for the Environment. Y. Z. was supported by the Schmidt Science Fellowship, in partnership with the Rhodes Trust. We are grateful to members of the Dionne and Boehm labs, and particularly thank Dr. Parivash Moradifar and Alan Dai for support in this work. Part of this work was performed at the Stanford Nano Shared Facilities (SNSF), supported by the National Science Foundation under award ECCS-2026822. Some of this work was performed at the Stanford-SLAC Cryo-EM Facilities, supported by Stanford University, SLAC and the National Institutes of Health S10 Instrumentation Programs. From these facilities, we would like to thank Dr. Elizabeth Montabana, Dr. Magda Zaoralova, and Dr. Bharti Singal for providing training and support in cryoEM imaging. Nanoparticle concentrations were measured by Guangchao Li at the Stanford Environmental Measurements Facility.

