## Supplementary material for "Bacterial Wastewater-Based Epidemiology Using Surface-Enhanced Raman Spectroscopy and Machine Learning": Fig. S6

**Figure S6**

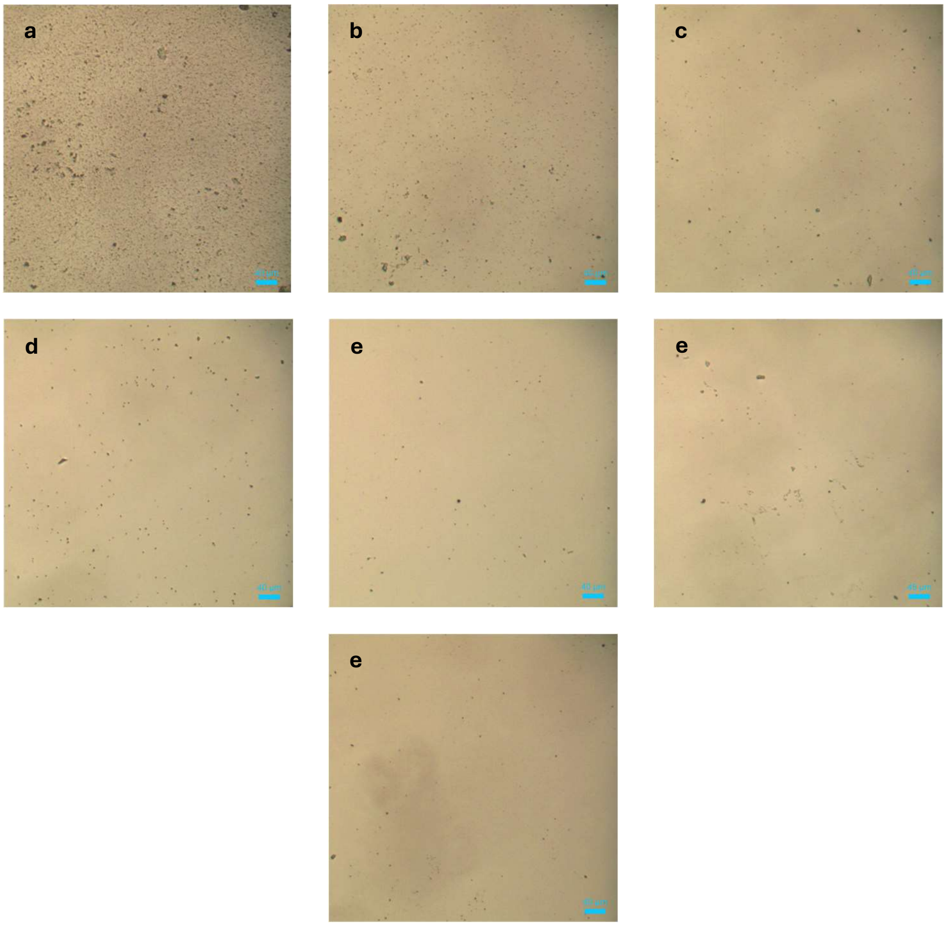


Liquid wells containing *S. aureus* in formalin at concentrations of (a) $10^9$ cells/mL, (b) $10^8$ cells/mL, (c) $10^7$ cells/mL, (d) $10^6$ cells/mL, (e) $10^5$ cells/mL, (f) $10^4$ cells/mL, and (g) 0 cells/mL. Particles visible in sample with no bacteria re likely dust and debris from tape.
