## Supplementary material for "Bacterial Wastewater-Based Epidemiology Using Surface-Enhanced Raman Spectroscopy and Machine Learning": Fig. S5

**Figure S5**

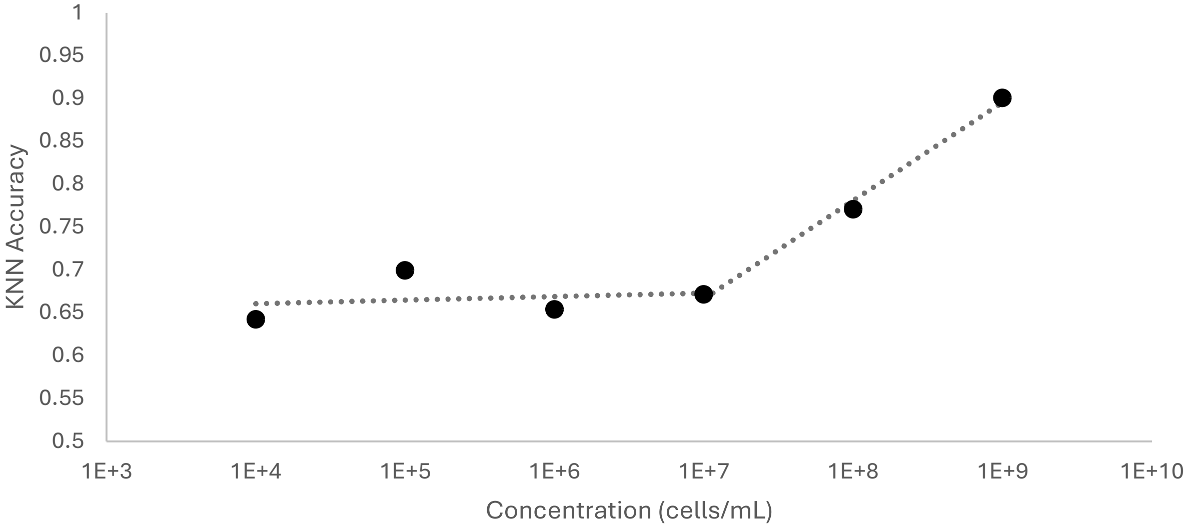


KNN Classification accuracy plotted against bacterial concentration. Linear regressions were used to identify linear relationships between accuracy and log(concentration) in concentration ranges from $10^4$-$10^7$ cells/mL and from $10^7$-$10^9$ cells/mL. This inflection point of $10^7$ cells/mL was selected because it minimized the root mean squared error of the linear fits, compared to regressions with other inflection points.
