## Supplementary material for "Bacterial Wastewater-Based Epidemiology Using Surface-Enhanced Raman Spectroscopy and Machine Learning": Fig. S4

**Figure S4**

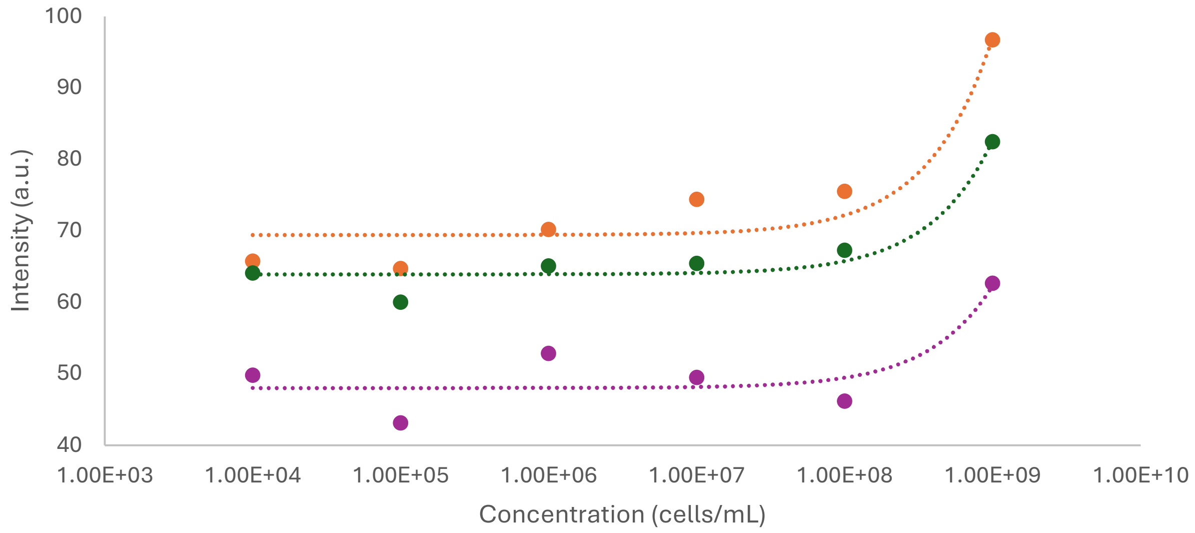


Average peak intensities for spectra with bacteria at each concentration at 851 $cm^{-1}$ (orange), 1040 $cm^{-1}$ (green), and 1,599 $cm^{-1}$ (purple). These values display a positive correlation between bacterial concentration and peak intensity, which can be fit to linear regressions, represented by dotted lines.
