## Supplementary material for "Bacterial Wastewater-Based Epidemiology Using Surface-Enhanced Raman Spectroscopy and Machine Learning": Fig. S3

**Figure S3**

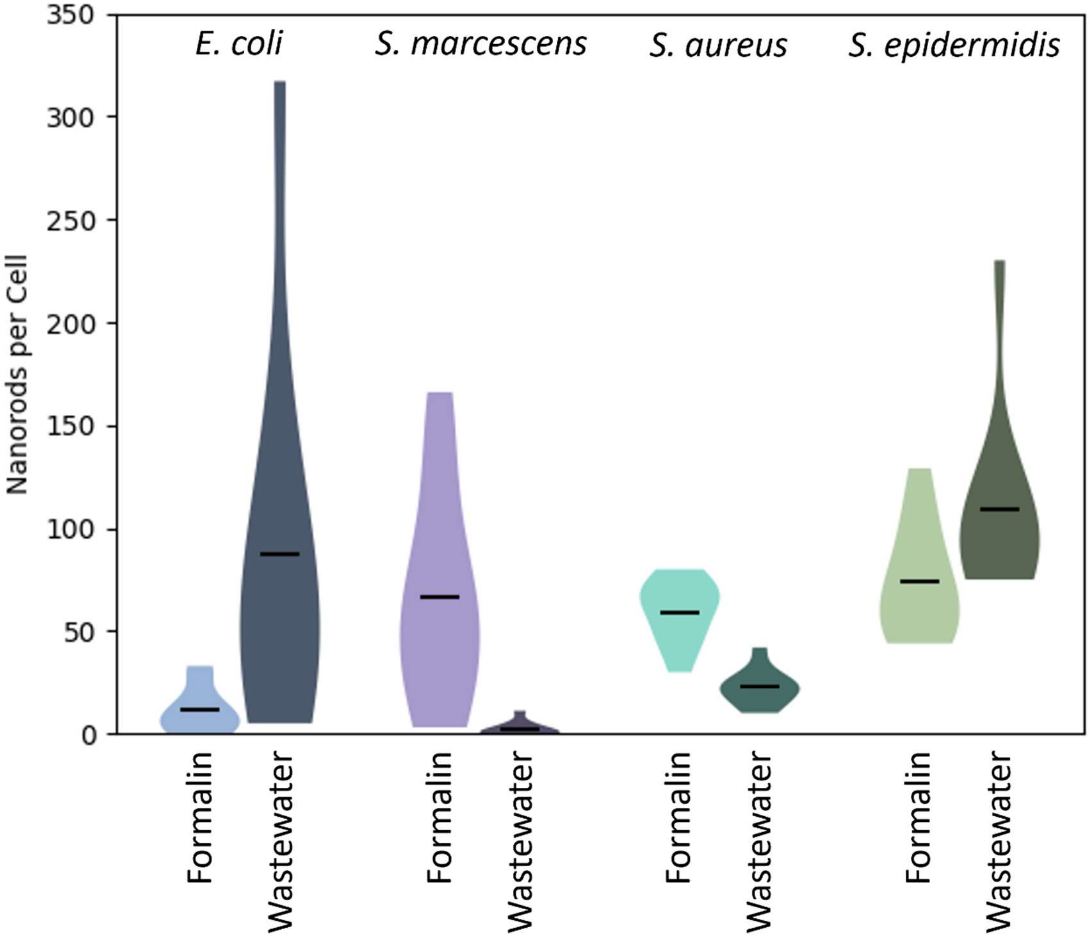


Violin plot displaying number of AuNRs bound to each species in formalin and wastewater. Each distribution consists of 10 transmission cryoelectron micrographs, each of a single cell. For each micrograph, the number of AuNRs in contact with the cell was counted by hand. The black line indicates the median of each distribution. For all species except *S. marcescens*, AuNRs all bacteria in wastewater had AuNRs bound to their surface. For *S. marcescens*, while some bacteria did not have AuNRs on their surface, a majority of them still did.
