## Supplementary material for "Bacterial Wastewater-Based Epidemiology Using Surface-Enhanced Raman Spectroscopy and Machine Learning": Fig. S2

**Figure S2**

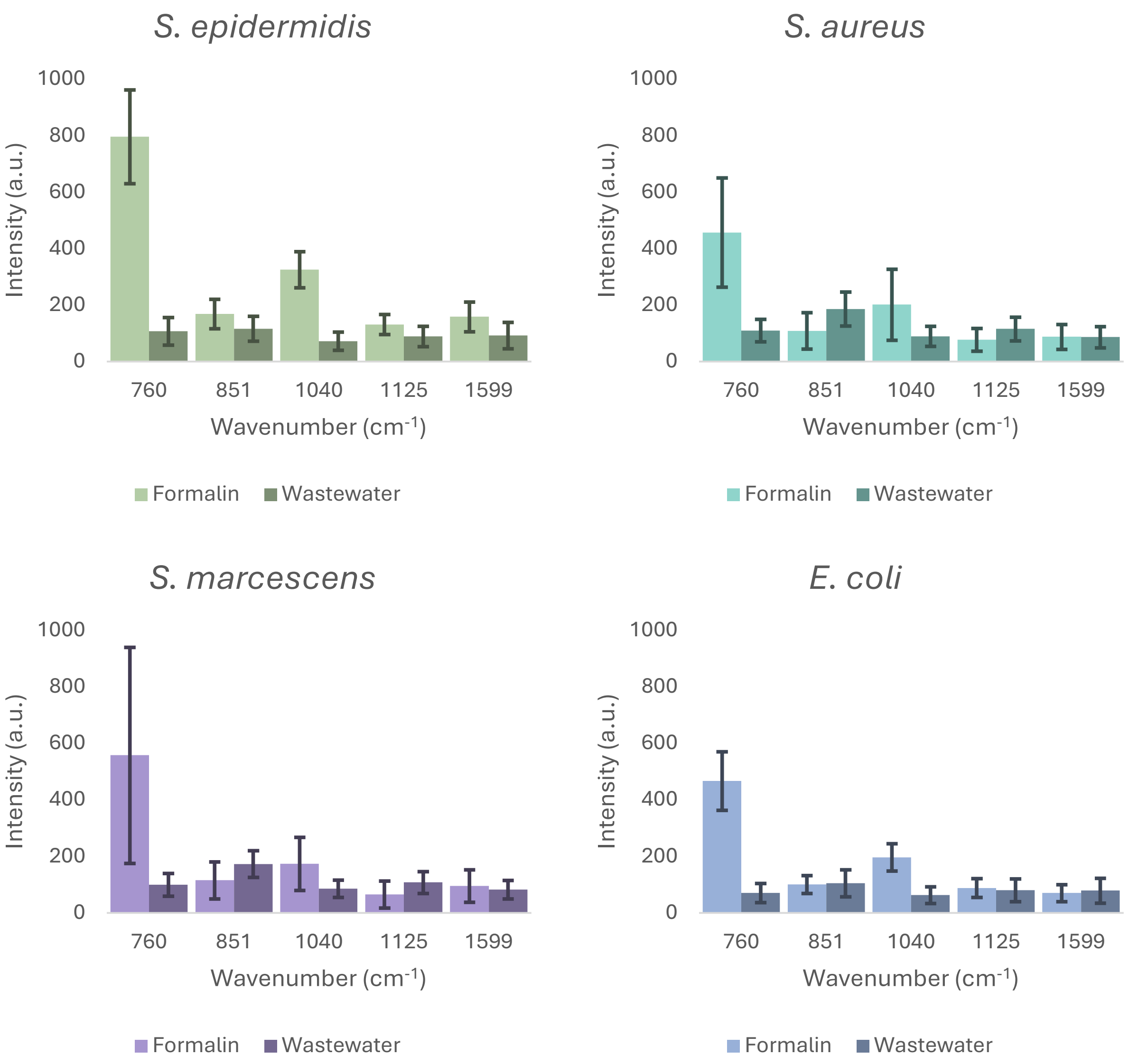


Intensities of selected Raman peaks for all four species in formalin and filtered wastewater with an AuNR concentration of 150 µg/mL. For all four species, intensities in wastewater display a statistically significant ($p<0.01$) decrease compared to formalin at the 760 $cm^{-1}$ and 1,040 $cm^{-1}$ peaks, while no statistically significant difference in intensity is observed for the other three peaks.
