## Supplementary material for "Bacterial Wastewater-Based Epidemiology Using Surface-Enhanced Raman Spectroscopy and Machine Learning": Fig. S1

**Figure S1**

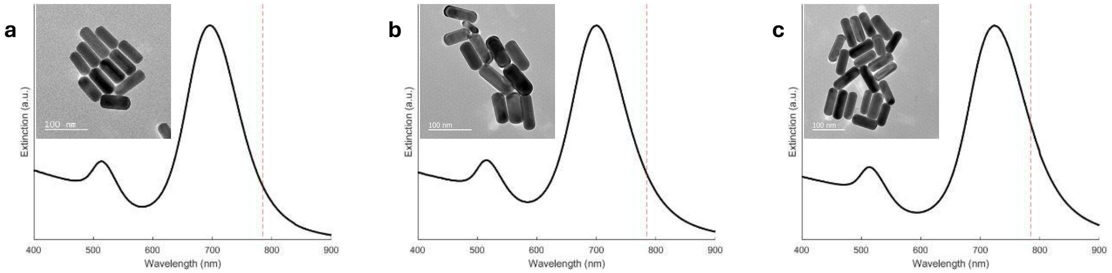


Extinction spectra and transmission electron micrographs of AuNRs used in experiments described in (a) Figures 1, 3, and 4; (b) Figure 5; and (c) Figure 2. All show week transverse resonances at 520 nm and strong longitudinal resonances near 700 nm.
